# Ginkgolic acid attenuates *echinococcus granulosus* infection-induced hepatic fibrosis by inhibiting Smad4 SUMOylation

**DOI:** 10.1101/2025.08.22.671734

**Authors:** Qiuyue Chen, Dan Dong, Xueting Yu, Xinyu Jiang, Huijiao Jiang, Jun Hou, Lianghai Wang, Junying Xu, Xiangwei Wu, Xueling Chen

**Author notes:** These authors contributed equally to this work (Qiuyue Chen, Dan Dong). Correspondence (Xiangwei Wu); (Xueling Chen).

## Abstract

**Background:** The SUMOylation modification is closely linked to the progression of fibrotic diseases, yet its role in hepatic fibrosis associated with cystic echinococcosis (CE) remains unclear. This study aimed to investigate the function of SUMOylation in CE-related hepatic fibrosis and evaluate the anti-fibrotic effects and mechanisms of ginkgolic acid (GA) via regulation of the SUMOylation pathway.

**Methodology:** Peri-lesional (PL) and adjacent normal (AN) liver tissues from CE patients were collected to examine histopathology and SUMO pathway proteins. A CE-infected mouse model was established and treated with GA to assess cyst burden, serum TGF-β1 levels, hepatic fibrosis markers, and SUMO-related proteins. *In vitro*, macrophages and hepatic stellate cells (HSCs, LX-2 line) were stimulated with *Echinococcus granulosus* cyst fluid (EgCF) or TGF-β1 to evaluate GA’s effects on macrophage polarization (CD206/CD86), HSC activation (α-SMA/PCNA), Smad4 SUMOylation, and nuclear translocation. Macrophage-HSC crosstalk was investigated via conditioned medium co-culture assays.

**Result:** Fibrosis was exacerbated in peri-lesional liver tissues of CE patients, accompanied by SUMO pathway activation. GA significantly alleviated hepatic fibrosis in CE mice and reversed SUMO pathway dysregulation. Mechanistically, GA inhibited EgCF-induced pro-fibrotic M2 macrophage polarization and blocked Smad4 SUMOylation and nuclear translocation by modulating SUMOylation. Furthermore, GA directly suppressed HSC activation and bidirectionally disrupted the pro-fibrotic crosstalk between macrophages and HSCs under EgCF stimulation, ultimately alleviating fibrosis.

**Conclusion:** This study reveals the critical role of SUMOylation modification in CE-associated hepatic fibrosis and elucidates a novel anti-fibrotic mechanism whereby GA targets the SUMOylation-Smad4 axis to regulate the immune microenvironment.

**Author summary:** This study demonstrates aberrant activation of the SUMOylation pathway during hepatic fibrosis progression in cystic echinococcosis (CE), characterized by upregulated SUMO1/Ubc9 expression and downregulated SENP1 in peri-cystic liver tissue. Ginkgolic acid (GA) intervention significantly attenuated CE-associated liver fibrosis in mice, evidenced by reduced cyst volume, decreased TGF-β1 levels, and suppressed expression of fibrotic markers (α-SMA/COL1A1). GA concurrently reversed dysregulated SUMO pathway protein expression. Mechanistically, GA upregulates the deSUMOylating enzyme SENP1, thereby inhibiting Echinococcus granulosus cyst fluid (EgCF)-induced SUMOylation and nuclear translocation of Smad4. This blockade impedes macrophage polarization toward the pro-fibrotic M2 phenotype (CD206↓) and suppresses hepatic stellate cell (HSC) activation (α-SMA/PCNA↓). Furthermore, GA disrupts the pro-fibrotic bidirectional crosstalk between HSCs and macrophages. Collectively, these findings indicate that GA ameliorates CE-induced hepatic fibrosis by targeting the SUMO-Smad4 axis to modulate the immune microenvironment, providing a novel therapeutic strategy.

## Introduction

Cystic echinococcosis(CE), caused by *Echinococcus granulosus* sensu lato, is a zoonotic parasitic disease that seriously endangers public safety and high endemicity in pastoral areas such as western China, central Asia [1, 2]. The WHO recognizes echinococcosis as one of 20 neglected tropical diseases and prioritizes it for targeted control measures [3]. Humans CE is a slow-growing cyst in the liver that induces inflammation, liver fibrosis and other immunopathological reactions. Surgery and drugs such as albendazole are commonly used in clinical treatment [4]. However, these traditional methods are difficult to avoid sac effusion or disease recurrence.

Therefore, it is urgent to find new CE treatment drugs. Ginkgo acid (GA) is a kind of small molecular phenolic organic compound, and many of its functions are closely related to SUMOylation [5]. In recent years, it has been found that it has many pharmacological activities such as anti-tumor, anti-virus, anti-inflammatory and anti-fibrotic [6]. However, little is known about the effect of GA on CE liver fibrosis, which needs further study.

SUMOylation is an important post-translational protein modification, which is a three-stage enzymatic cascade of SUMO molecules bind to target proteins under the catalysis of E1 activating enzyme, E2 binding enzyme (Ubc9) and E3 ligase. In addition, this process can be reversed by SENP to form a dynamic balance of SUMOylation and deSUMOylation [7, 8]. SUMOylation modifications regulate a variety of cellular processes, including transcription, immune response, protein activity, and protein-protein interactions[8]. The dysfunction of its modification function is closely related to the occurrence and development of liver diseases. It was found that alcohol can induce CYP2E1-SUMOylation *in vitro* and *in vivo* [9, 11]. Hepatitis virus replication is related to SUMO1 modification [10, 11]. Down-knocking of Ubc9 inhibits SUMOylation, prevents the development of liver fibrosis and inhibits TGF-β / Smad signaling [11]. However, whether SUMOylation is involved in hepatic fibrosis in CE is unclear. Recently, it has been reported that GA as a SUMO1 inhibitor, can promotes inflammatory responses by inhibiting the SUMOylation of NF-κB p65 in macrophages [12], also can anti-hepatic fibrosis by inhibiting the expression of SUMO1-activating enzyme subunit 1 (SAE1) [13], which suggests its may be important in CE liver fibrosis.

In the early stage of CE, M1 macrophages are recruited to the liver to release cytokines that combat parasitic infection. In the middle and late stages of infaection, M2 macrophages play an anti-inflammatory or tissue repair role, while activated hepatic stellate cells (HSCs) secrete abundant extracellular matrix (ECM) proteins, promoting the development of chronic fibroinflammatory state and pathological fibrosis [14, 15]. The TGF-β / Smad signaling pathway plays a crucial role in the development of liver fibrosis caused by CE [16]. Smad4 serves as a central mediator in TGF-β / Smad signal transduction and shuttles between the cytoplasm and nucleus. SUMOylation of Smad4 protein occurs, forming a complex with SUMO1 [17]. At present, it is unknown whether Smad4 SUMOylation is involved in CE liver fibrosis and whether it is related to the protective effect of ginkgo acid on CE liver fibrosis. Therefore, we first conducted the pathological relevance of SUMOylation in CE-associated hepatic fibrosis by clinical specimens. Then, evaluated the therapeutic efficacy of GA on murine cystic echinococcosis models and macrophage cultures. Finally, through co-immunoprecipitation and nucleocytoplasmic fractionation revealed that GA attenuated M2 polarization via Smad4 SUMOylation, and disrupted macrophage-HSC crosstalk. These findings provides a new idea for the prevention and treatment of CE.

## Methods

### Chemicals and reagents

Ginkgolic acid (MCE, HY-N0077), β-mercaptoethanol (Gibco, USA), phorbol 12-myristate 13-acetate (PMA) (MultiSciences, CS0001), Lipopolysaccharides (LPS) (Solarbio, L8880), recombinant human TGF-β1 protein (Active) (Abcam, ab50036). All reagents were reconstituted and used according to the manufacturers’ instructions. Details of the antibodies used are provided in Supplementary Table 1.

### Ethics statement

The study strictly followed the relevant ethical guidelines. All clinical samples were obtained after written and signed informed consent, which was approved by the Ethics Committee of Science and Technology of the First Affiliated Hospital of Shihezi University (approval number: KJ2025-003-01). The experimental procedures involving animals followed the principles of animal welfare and were reviewed and approved by the Bioethics Committee of Shihezi University (approval number: A2024-427).

### Animal experiments

Female C57BL/6 mice were purchased from SpePharm (Beijing) Biotechnology Co., Ltd. (Laboratory Animal Production License: SCK (Jing) 2019-0010). After one week of acclimatization under controlled conditions (22°C - 24°C, 40%-60% humidity, 12 / 12-hour light/dark cycle), the experiment commenced. Thirty-two mice were randomly divided into four groups (n=8): vehicle control group: Intraperitoneal (i.p.) injection of an equivalent volume of saline; GA safety group: i.p. injection of GA (25 mg/kg/day); CE group: *Echinococcus granulosus* (hydatid disease) infection mode; CE+GA Group: *E. granulosus* infection combined with i.p. injection of GA (25 mg/kg/day) [5]. The mice in CE group and CE+GA group were anesthetized, and inoculated with protoscoleces (3000 per mouse) by subcapsular injection, as described previously [18]. Eight weeks after infection, the GA and CE+GA groups were intraperitoneally injected with GA (25 mg/kg/d), and the solvent control group was injected with the same volume of normal saline. After 4 weeks of continuous administration, all mice were euthanized, and serum and target organ tissues were collected for subsequent experiments.

### Histopathological, Immunohistochemical, and Immunofluorescence Staining

Clinical and animal tissue samples were routinely dehydrated, embedded in paraffin, and sectioned after fixation with 4% paraformaldehyde for 24 h. HE staining was used to evaluate histopathological damage, and Masson and Sirius Red staining were used to evaluate the degree of liver fibrosis. For immunohistochemical (IHC) analysis, based on the mouse/rabbit hyper-sensitivity polymer detection system (PV-8000, Zhongshan Bridge), sections were dewaxed hydrated, antigen repaired, incubated overnight at 4 ℃ with primary antibodies specific for the target protein (see Table 1 in the Supplementary Material for details), incubated with hyper-sensitive enzyme-labeled secondary antibodies, and examined by DAPI sealed microscopy. Image software was used for quantitative evaluation. Immunofluorescence staining analysis: the sections were deparaffinized and hydrated, followed by antigen repair and endogenous peroxidase inactivation. Dual-color labeling of F4 / 80 primary antibody / iF555-Tyramide secondary antibody and SUMO1 primary antibody / iF488-Tyramide secondary antibody was performed in turn, and the slides were blocked with DAPI and imaged using fluorescence microscopy.

### Enzyme-linked immunosorbent assay (ELISA)

Mouse serum TGF-β1 levels were quantified using a mouse-specific TGF-β1 ELISA kit (JL12223-48T, JONLNBIO, China), while IL-13 and TGF-β1 levels in cell supernatants were measured using a Human IL-13 ELISA Kit (EK113, MULTISCIENCES, China) and a Human TGF-β1 ELISA Kit (EK981, MULTISCIENCES, China), respectively; all assays were performed in strict accordance with the manufacturers’ instructions.

### Cell experiments

Human hepatic stellate cells (LX-2), human monocytic cells (THP-1), and mouse monocyte-macrophage cells (RAW 264.7) were obtained from the Cell Bank of the Chinese Academy of Sciences. LX-2 and RAW 264.7 cells were cultured in DMEM medium (Gibco, China) supplemented with 10% fetal bovine serum (FBS) (Procell, China) and 1% penicillin/streptomycin (Solarbio, China). THP-1 cells were cultured in suspension in 1640 medium containing 10% FBS, 1% penicillin/streptomycin, and 0.05 mM β-mercaptoethanol. Prior to use, THP-1 cells were differentiated into adherent macrophages by treatment with 100 nM phorbol 12-myristate 13-acetate (PMA) for 24 h. All cells were maintained and passaged in a humidified incubator at 37℃ with 5% CO₂. RAW 264.7 cells or PMA-induced differentiated THP-1 cells were treated with LPS, LPS+EgCF, or LPS+EgCF+GA for 12 h before WB analysis. LX-2 cells were treated with TGF-β1 (10 ng/mL) or EgCF (1 mg/mL) in combination with different concentrations of GA (10 nM, 20 nM) for 12 h before WB analysis. Cell interaction experiment: LX-2 cells were treated with the supernatant of LPS stimulated THP-1 cells (CM1) or LPS+EgCF stimulated THP-1 cells (CM2), followed by GA treatment for 12 h. Second, THP-1 cells were treated with EgCF-stimulated LX-2 cell supernatant (CM) followed by GA treatment for 12 h, and the cells were collected for WB analysis.

### Western blot analyses (WB)

After tissue or cell samples were lysed with RIPA lysate containing protease inhibitors, the supernatants were collected by centrifugation at 12,000 ×g for 15 min. Protein concentration was determined by the BCA method, and equal amounts of adjusted proteins were separated by 10% SDS-PAGE electrophoresis and transferred to PVDF membrane under constant pressure. After blocking with BSA for 2 h, membranes were incubated with the corresponding primary antibody overnight at 4°C and with the secondary antibody for 2 h at room temperature. ECL chemiluminescence reagent was used for visualization, and Image J software was used to quantify the gray value of the target band. Statistical analyses were performed by GraphPad Prism 8.

### Immunofluorescence (IF)

The THP-1 cells in different groups were successively fixed with 4% paraformaldehyde for 30 min, permeabilized with 0.3% Triton X-100 for 10 min, and blocked with 5% BSA for 30 min. Then, they were incubated with the primary antibodies SUMO1 and Smad4 at 4℃ overnight. Subsequently, the cells were incubated at room temperature in the dark with the fluorescently labeled secondary antibodies for 2 h. After DAPI staining to re-dye the cell nuclei, images were captured using a fluorescence microscope (NIKON ECLIPSE C1, 100×). The detailed information of the antibodies is shown in Table S1.

### Co-immunoprecipitation (Co-IP)

Following cell lysis on ice using RIPA lysis buffer containing a protease inhibitor cocktail, anti-Smad4 antibody and control antibodies were added to the lysates and incubated overnight to form protein-antibody complexes, while 10% of the lysate was retained as a positive control; subsequently, protein A/G magnetic beads (RM02915, ABclonal, China) were added to the protein-antibody complexes and incubated at 4 °C for 4 h to facilitate bead-antibody-protein complex binding; the beads were then centrifuged, the supernatant carefully aspirated, and the bead-protein complexes retained; the bead-protein complexes were then washed repeatedly to remove non-specifically bound material; finally, the washed beads were resuspended in 1× SDS loading buffer and boiled at 100 °C for 10 min; the resulting samples were then subjected to WB analysis according to standard protocols to detect the captured target proteins and assess the impact of each experimental factor on protein-protein interactions.

### Nuclear and Cytoplasmic Extraction

Nuclear and cytoplasmic proteins were isolated from THP-1 cells using a Nuclear and Cytoplasmic Protein Extraction Kit (Beyotime, China) according to the manufacturer’s instructions, followed by WB to assess the expression levels of the target proteins in the nuclear and cytoplasmic fractions, where Histone H3 and GAPDH were used as loading controls for the nucleus and cytoplasm, respectively, and Smad4 served as the target protein.

### Statistical analysis

Data analysis and processing were performed using GraphPad Prism 8.0 software. The statistical significance between two groups was determined using an unpaired t-test, and comparisons among three or more groups were performed using one-way ANOVA. Data are reported as mean ± SD, with statistical significance set at p < 0.05: *p < 0.05, **p < 0.01, ***p < 0.001.

## Results

### Altered expression of SUMO proteins in the lesion microenvironment of hepatic CE patients

A growing number of studies have shown that SUMOylation is closely related to the progression of fibrosis related diseases [6, 11]. Currently, It is unclear whether CE liver fibrosis is associated with SUMOylation. First, liver paraffin samples from hepatic CE patients were collected and histopathological staining were performed to confirm fibrosis. Liver tissue near the CE lesion was called peri-lesion (PL) specimen, while tissue about 5 cm away was termed adjacent normal (AN) specimen [19]. Compared to the AN liver, the PL liver demonstrated a substantial increase in inflammatory cell infiltration and a higher density of collagen fiber deposition. The expression levels of a-SMA, COL1A1, SUMO1, and Ubc9 were significantly upregulated, while SENP1 expression was markedly downregulated (Fig 1). The above results preliminarily indicated that CE liver fibrosis is associated with SUMO proteins.

**Fig 1.**
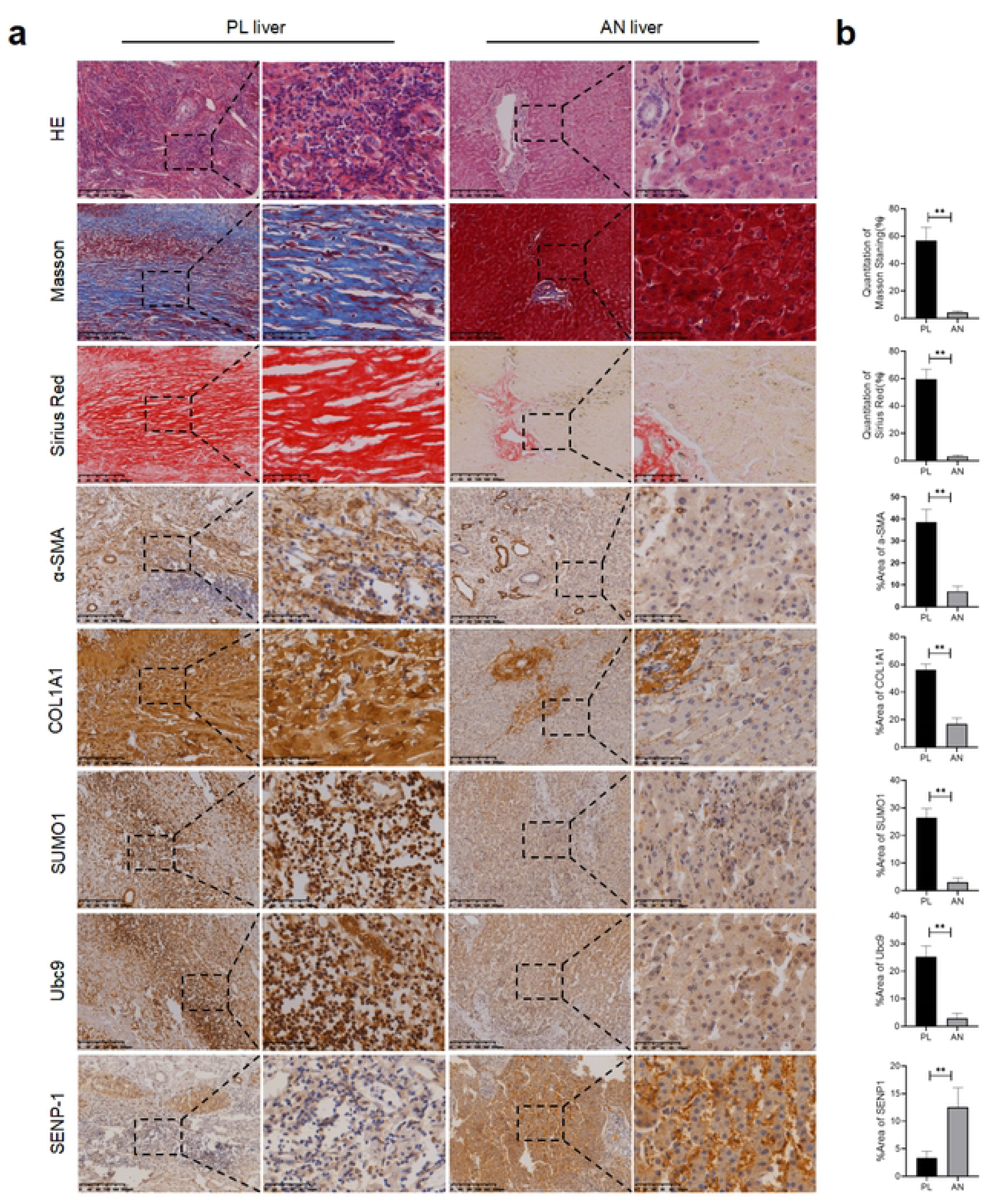
Differential expression of SUMO proteins in the lesion microenvironment. of hepatic CE patients. (a) Representative H&E, Masson, Sirius red and immunohistochemical (IHC) staining of a-SMA, COL1A1, SUMO1, Ubc9 and SENP1 in peri-lesion (PL) liver and adjacent normal (AN) liver obtained from hepatic CE patients. (b) Quantitative analysis of positively stained areas for Masson staining, Sirius red and α-SMA, COL1A1, SUMO1, Ubc9, SENP1 expression. *p < 0.05, **p < 0.01 and ***p < 0.001.

### GA modulated SUMO1, Ubc9, and SENP1 expression in Echinococcus granulosus-infected liver fibrosis mouse model

Studies have shown that GA can be used as SUMOylation inhibitors play a role. We hypothesized that GA might attenuate CE liver fibrosis by regulating the expression of SUMO proteins. A model of CE was established in female C57BL/6 mice via injection of protoscoleces (PSCs) to the liver capsule. From the 8th week to the day before sacrifice (total 28 days), mice were Intraperitoneal injectioned daily with GA (GA, 25 mg/ kg) (Fig 2a). Compared with the CE group, the CE+GA group of the liver cysts volume were significantly reduced (Fig 2b and 2c). And the serum TGF-β1 levels were markedly decreased (Fig 2d). Masson staining, Sirius red staining and a-SMA, COL1A1 IHC staining results show that the expression of a-SMA and COL1A1 increased, the HSCs were activated, ECM proteins accumulated, large amounts of collagen were deposited, and accompanied by granulomas, calcification in the PL tissue of the liver in the CE group. All of which were significantly reduced by GA treatment (Fig 2e and 2f). Besides, the histological HE staining examination showed that in the mice given GA (25 mg/kg, for 4 weeks), no obvious pathological changes were observed in their heart, liver, spleen, lung, and kidney tissues, and there was no significant difference compared with the normal control group. This indicates that GA has no obvious systemic toxicity at this dose and is safe (Fig S1). Then, we further studied the relationship between GA treatment of CE mice and SUMO proteins through IHC staining and WB. The results show that the expression of SUMO1 and Ubc9 were increased, while the expression of SENP1 protein were decreased in the CE group. The expression changes of these proteins can be reversed by GA action (Fig 2e and 2g).

**Fig 2.**
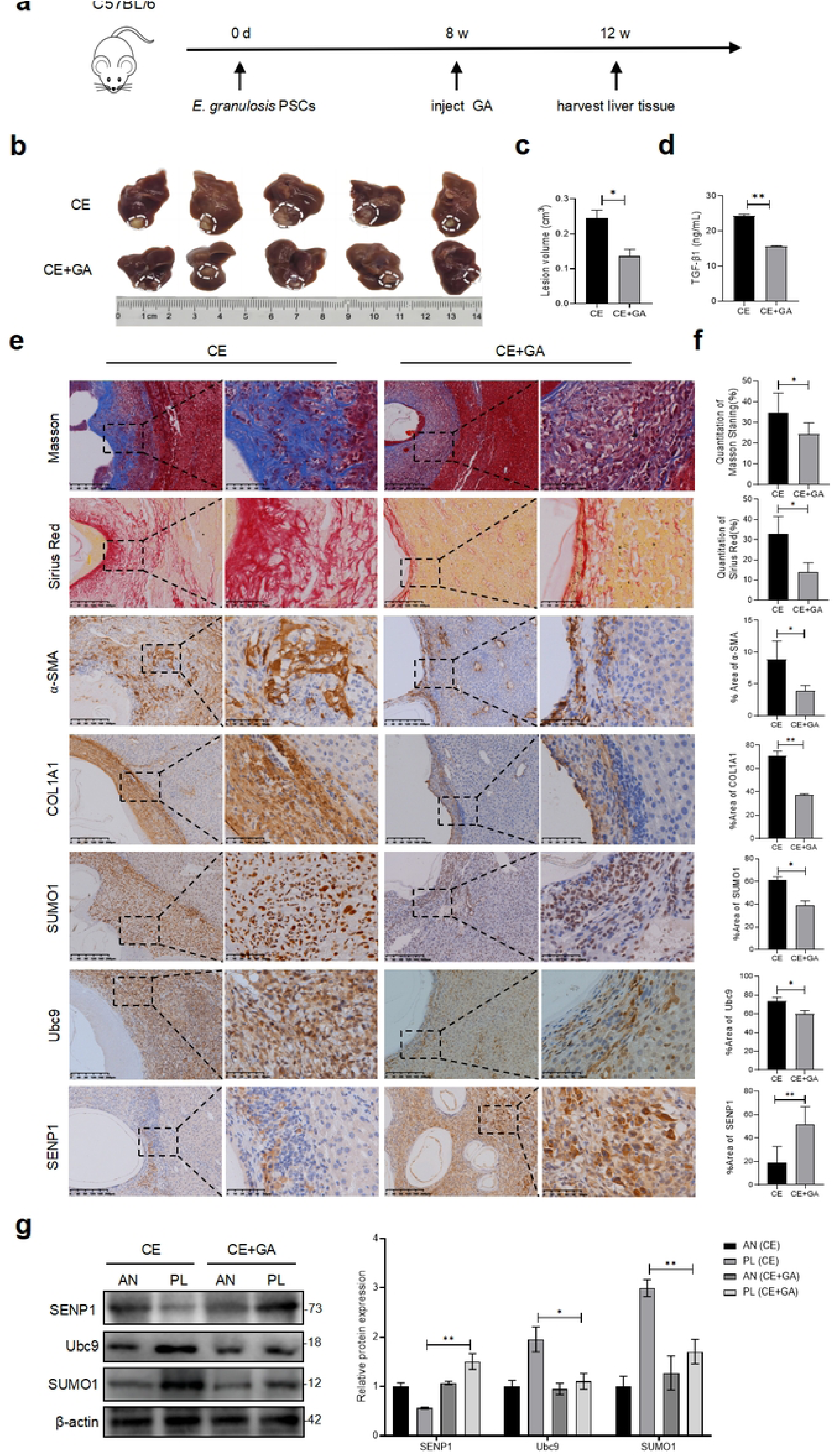
GA modulated key SUMO proteins in vivo. (a) Mice injected with PSCs under the liver capsule were administered GA intraperitoneally at 8 weeks, and the experiment concluded at 12 weeks. (b) Gross images of the livers of mice in each group. (c) Lesion volume were calculated and compared among two groups. (d) Serum TGF-β1 levels of the two groups were measured by ELISA. (e) Representative Masson, Sirius red and IHC staining of a-SMA, COL1A1, SUMO1, Ubc9 and SENP1 in peri-lesion (PL) liver from two groups. (f) Quantitative analysis of positively stained areas for Masson staining, Sirius red and α-SMA, COL1A1, SUMO1, Ubc9, SENP1 expression. (g) Western blot (WB) analysis of the protein expression levels of SUMO1, Ubc9, SENP1 in peri-lesion (PL) or adjacent normal (AN) liver from two groups mice. *p < 0.05, **p < 0.01 and ***p < 0.001.

### GA Downregulated EgCF-induced inflammatory responses in macrophages

The aforementioned findings demonstrated that the expression levels of SUMO1, Ubc9 were elevated in parallel with the progression of CE-associated hepatic fibrosis, whereas GA administration significantly upregulated the expression of SENP1 in mice models, concomitantly attenuating the severity of liver fibrosis. Notably, clinical data from CE patients and experimental mice models of CE revealed a positive correlation between M2 macrophage polarization markers (CD206, Arg-1) and fibrotic progression, as documented in our prior research [20, 21]. Based on these observations, we hypothesized that GA may exert anti-fibrotic effects via modulation of macrophage-mediated inflammatory responses. As illustrated in Figure 3, subsequent investigations were conducted utilizing both *in vivo* murine models and in vitro macrophage systems. Tissue immunofluorescence analysis of hepatic tissues demonstrated that GA treatment markedly reduced the infiltration of SUMO1-positive / F4/80-positive macrophages (Fig 3a). Furthermore, *in vitro* inflammatory models of THP-1 and RAW264.7 macrophages induced by Echinococcus granulosus cystic fluid (EgCF) were established to evaluate GA’s therapeutic potential. Western blot analysis revealed that EgCF stimulation significantly upregulated M2 polarization marker CD206 while suppressing M1 marker CD86 expression in THP-1 cells, whereas GA intervention potently inhibited its expression level (Fig 3b). A consistent inhibitory trend was observed in RAW264.7 macrophages following analogous treatments (Fig 3c). Collectively, these data substantiate that GA effectively suppresses EgCF-induced M2 macrophage polarization.

**Fig 3.**
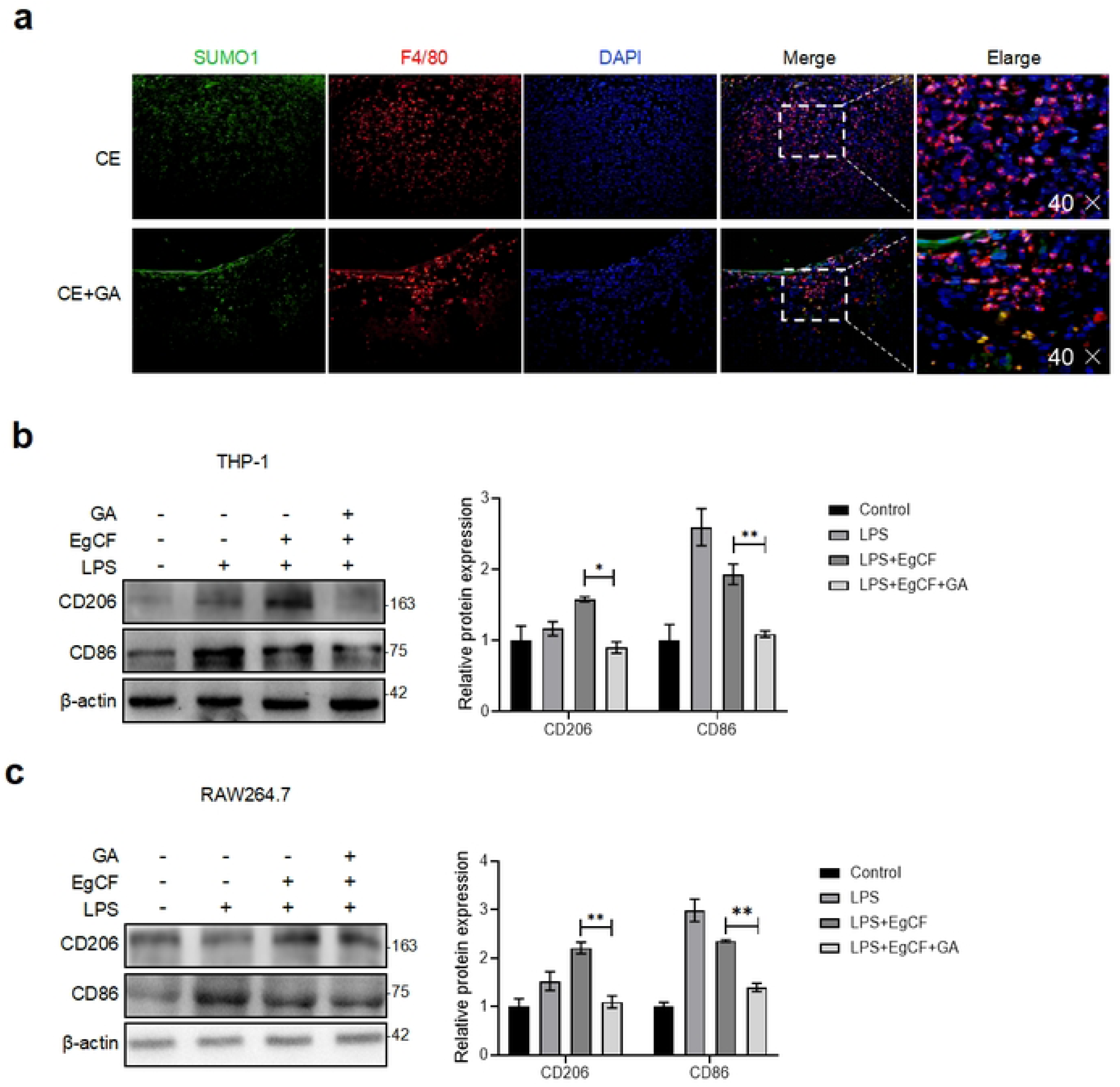
GA inhibited macrophage polarization induced by EgCF. (a) Immunofluorescence (IF) staining for SUMO1 (green) and F4/80 (red) were performed on Paraffin liver tissue sections, and the nuclei were stained with DAPI (blue), magnification: 20 × and 40 ×. RAW 264.7 cells and THP-1 cells were pretreated with LPS (1 μg/mL) and then with EgCF (1 mg/mL) and GA (10 nM) for 12 h, CD206 and CD86 protein expression levels in RAW 264.7 cells (b) and THP-1 cells (c) were detected by WB. *p < 0.05, **p < 0.01 and ***p < 0.001.

### GA inhibited EgCF induced Smad4 SUMOylation

Targeted modulation of macrophage polarization serves as a critical therapeutic strategy for hepatic fibrosis [22]. Based on the regulatory mechanism of SUMOylation in macrophage inflammatory response pathways [23], and considering the established role of GA in modulating TGF-β1/Smad4 signaling in fibrotic diseases, we hypothesize that GA may regulate macrophage polarization through intervention in Smad4 SUMOylation. Firstly, total protein was extracted from the liver tissues of mice in each group. The WB results showed that, compared with the CE group, the protein expressions of SUMO1 and Smad4 decreased after GA treatment; the CO-IP results further indicated that GA inhibited the CE-induced Smad4 SUMOylation modification (Fig 4a). *In vitro*, we established an EgCF-induced macrophage inflammation model. Western blot analysis demonstrated that compared with the EgCF+LPS-treated group, GA intervention significantly downregulated protein expression of SUMO1, UBC9, and Smad4 while upregulating the deSUMOylating enzyme SENP1 (Fig 4b and 4c). CO-IP experiments further confirmed GA’s substantial inhibition of molecular interaction between Smad4 and SUMO1 (Fig 4d and 4e). Fluorescence microscopy observations revealed that GA treatment effectively reduced nuclear co-localization of Smad4 with SUMO1 (Fig 4f), and nuclear-cytoplasmic fractionation assays showed GA significantly suppressed Smad4 nuclear translocation (Fig 4g). These findings collectively indicate that GA promotes Smad4 deSUMOylation through SENP1 upregulation, thereby blocking nuclear translocation and subsequently inhibiting macrophage polarization towards the pro-fibrotic M2 phenotype.

**Fig 4.**
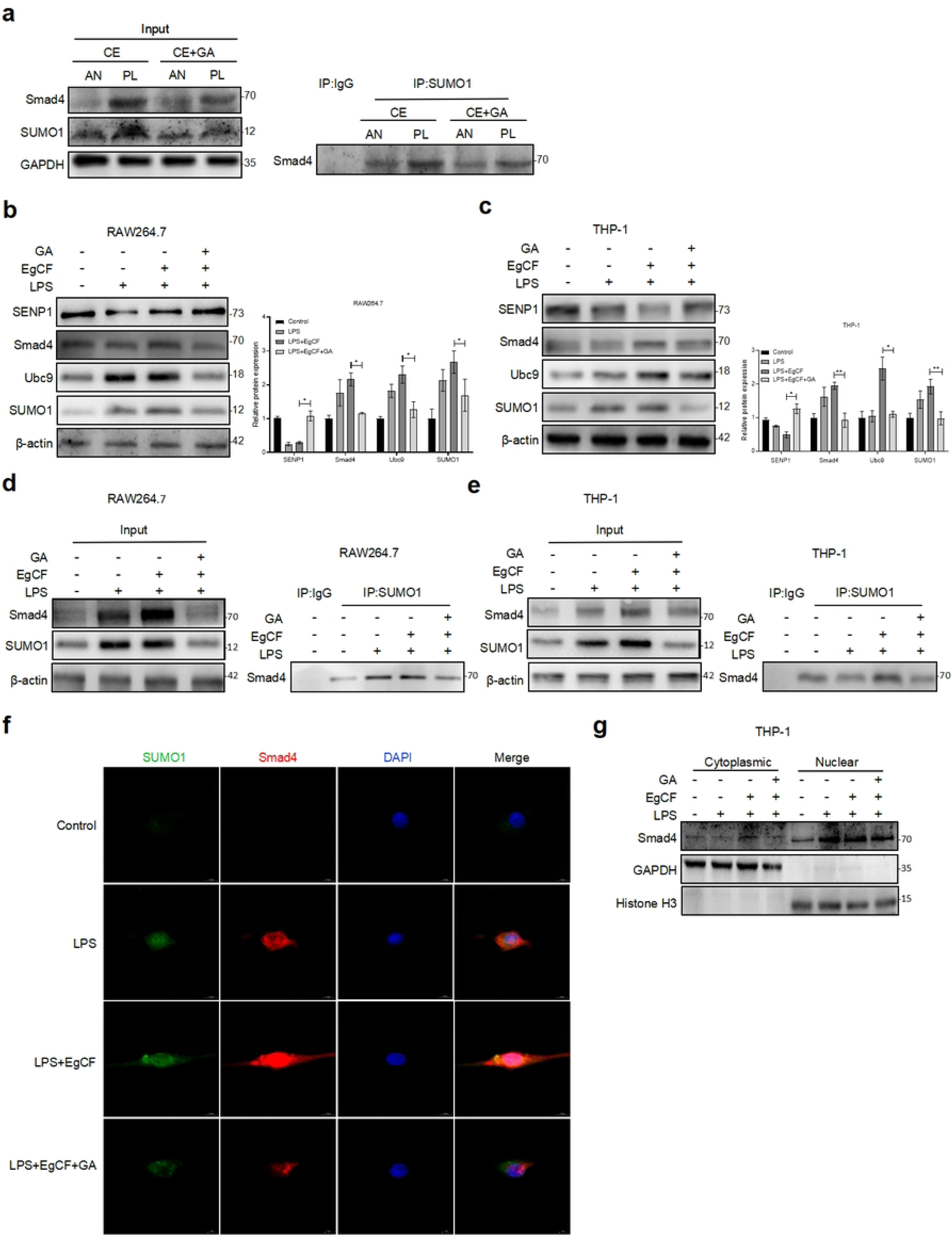
GA inhibited EgCF induced Smad4 SUMOylation. (a) SUMO1-Smad4 association in peri-lesion (PL) and adjacent normal (AN) liver from two groups mice: CO-IP with anti-SUMO1 antibody or Mouse IgG control, followed by Smad4-specific detection (WB), with Input controls showing both proteins. RAW 264.7 cells and THP-1 cells were pretreated with LPS (1 μg/mL) and then with EgCF (1 mg/mL) and GA (10 nM) for 12 h, SENP1, Smad4, UBC9 and SUMO1 protein expression levels in RAW 264.7 cells (b) and THP-1 cells (c) were detected by WB. SUMO1-Smad4 association in RAW 264.7 cells (d) and THP-1 cells (e) different groups: CO-IP with anti-SUMO1 antibody or Mouse IgG control, followed by Smad4-specific detection (WB), with Input controls showing both proteins. (f) Immunofluorescence double staining of SUMO1 (green) and Smad4 (red) in differentially treated THP-1 cell groups. Nuclei were counterstained with DAPI (blue), magnification: 100 ×. (g) Western blot for Smad4 expression in cytoplasmic and nuclear fractions of THP-1 cells treated with LPS (1 μg/mL) and then with EgCF (1 mg/mL) and GA (10 nM) for 12 h. *p < 0.05, **p < 0.01 and ***p < 0.001.

### GA inhibited the EgCF-induced HSCs activation and its crosstalk with macrophage polarization

To further investigate the involvement of GA in EgCF-induced hepatic stellate cell (HSC) activation and proliferation, we treated TGF-β1-or EgCF-stimulated LX-2 cells with varying GA concentrations (0, 10 nM, 20 nM) for 12 hours. Western blot analysis revealed that GA significantly suppressed the expression of proliferation marker PCNA and activation marker α-smooth muscle actin (α-SMA) in LX-2 cells (Fig 5a and 5b). Subsequently, LX-2 cells were co-cultured with THP-1 macrophage-derived conditioned medium (CM), where both CM1 (supernatant from LPS-stimulated THP-1 cells) and CM2 (supernatant from LPS+EgCF-stimulated THP-1 cells) were found to markedly upregulate PCNA and α-SMA levels in LX-2 cells. Notably, GA treatment specifically counteracted CM2-induced pro-fibrotic effects (Fig 5c). When THP-1 macrophages were co-cultured with LX-2-derived CM (supernatant from EgCF-stimulated LX-2 cells), GA significantly downregulated the expression of M2 polarization marker CD206 and macrophage activation marker CD86 in THP-1 cells (Fig 5d). Furthermore, GA effectively suppressed the secretion of IL-13 by LX-2 cells and TGF-β1 by THP-1 cells (Fig 5e). These findings collectively suggest that GA may exert its anti-fibrotic effects through bidirectional regulation of macrophage-HSC crosstalk, as evidenced by its capacity to attenuate both HSC activation/proliferation and macrophage polarization/activation in this reciprocal cellular communication system.

**Fig 5.**
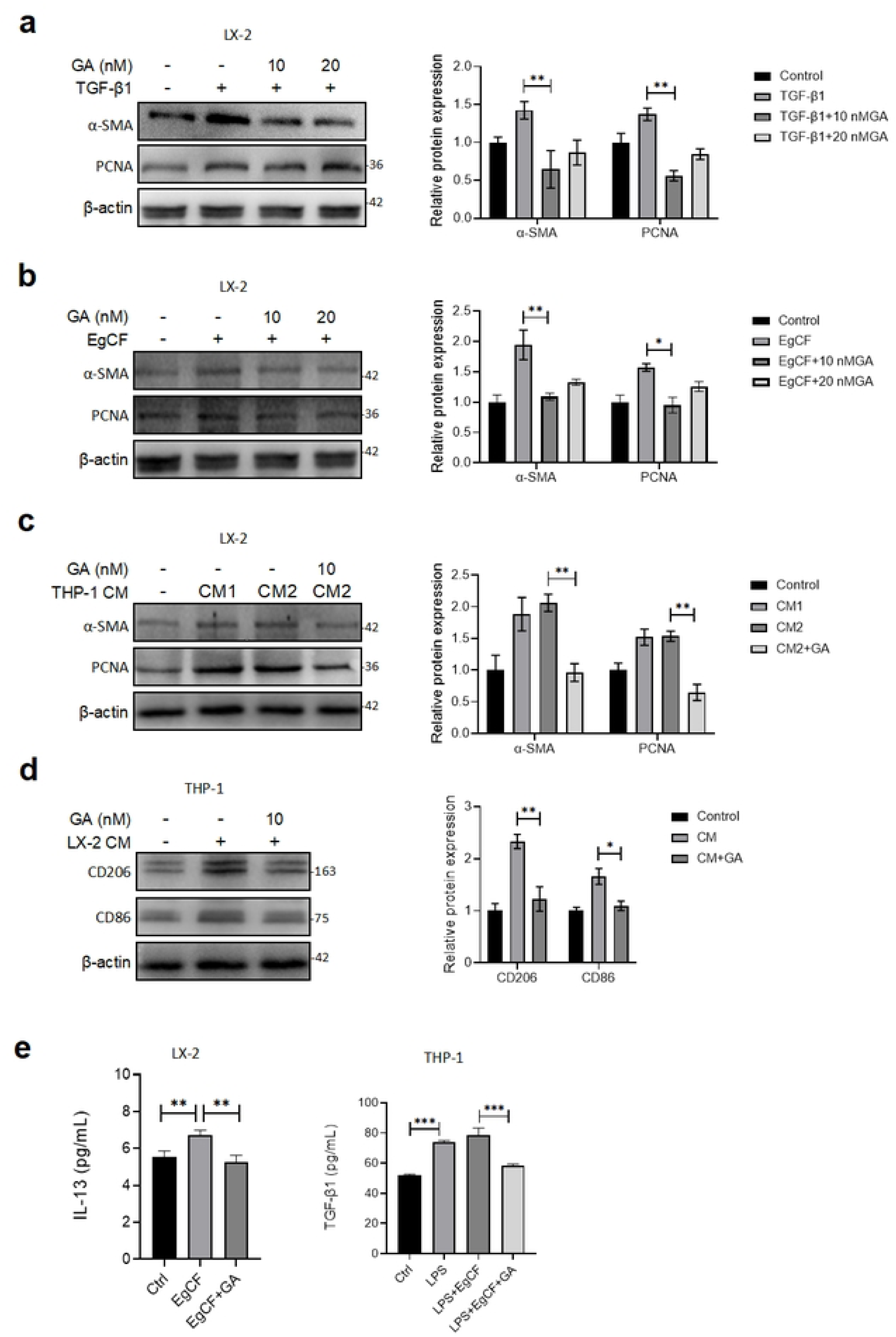
GA inhibited the EgCF-induced HSCs activation and its crosstalk with macrophage polarization. (a, b) LX-2 cells were treated with TGF-β1 (10 ng/mL) or EgCF (1 mg/mL) and GA (10 nM, 20 nM) for 12 h, the α-SMA and PCNA protein levels were analyzed by WB. (c) LX-2 cells were treated with conditioned medium (CM) from LPS-stimulated THP-1 cell ssupernatant (CM1) or LPS+EgCF-stimulated THP-1 cell ssupernatant (CM2) or CM2 and GA (10 nM) for 12 h, the α-SMA and PCNA protein levels were analyzed by WB. (d) THP-1 cells were treated with conditioned medium (CM) from EgCF-stimulated LX-2 cells ssupernatant for 12 h, the CD206 and CD86 protein levels were analyzed by WB. (e) Cell supernatant IL-13, TGF-β1 levels were measured by ELISA. *p < 0.05, **p < 0.01 and ***p < 0.001.

## 4. Discussion

Our research results show that GA ameliorates *E.granulosus* infection-mediated hepatic fibrosis by specifically inhibiting SUMOylation of Smad4 in macrophages through targeted modulation of the SUMOylation pathway, thereby hindering pathogenic macrophage-hepatic stellate cell crosstalk. This discovery not only reveals the central pathogenic driver role of the SUMO-Smad4 axis in parasite-induced fibrogenesis, but also provides a novel therapeutic target and candidate drug for anti-fibrotic treatment in hepatic echinococcosis.

Chronic Th2 inflammation triggered by *E.granulosus* persistently activates HSCs, causing them to transform into myofibroblasts, over-synthesize extracellular matrix (ECM), and abnormally deposit around the lesion, forming fibrous scars [24–25]. Recent studies demonstrate that SUMOylation and deSUMOylation play important roles in the process of heart, lung, liver and kidney fibrosis [5–6, 11, 26–28]. Our immunohistochemical results showed that in the lesion microenvironment of patients with cystic hepatic echinococcosis and infected mice, the expression of key molecules SUMO1, Ubc9 and SENP1 in the SUMOylation pathway was abnormal, suggesting that the imbalance of SUMOylation modification is a characteristic pathological event in the fibrotic microenvironment of hepatic echinococcosis. GA, as a natural SUMOylation inhibitor, exhibits therapeutic potential for counteracting organ fibrosis [5, 6]. Interestingly, GA significantly modulates expression levels of SUMO1, Ubc9, and SENP1 in *E. granulosus* infection-driven hepatic fibrosis models, further indicating the SUMOylation pathway as a key hub of the anti-fibrotic effect of GA. Our further research found that in the mouse model of *E.granulosus* infection, SUMO1 had significant spatial co-localization with the macrophage-specific marker F4/80. *In vitro* experiments further confirmed that EgCF stimulation could simultaneously up-regulate the expressions of SUMO1, E2 binding enzyme UBC9 and M2-type polarization marker CD206 in macrophages. However, GA intervention not only reduces the protein level of SUMO1, but also significantly weakens the co-localization signal of SUMO1 and F4/80, effectively reversing the M2-type hyperpolarization phenotype of macrophages induced by EgCF. Previous studies have shown that SUMOylation modification drives the core mechanism of the M2 polarization process in macrophages by dynamically regulating key signaling proteins, thereby mediating pathophysiological responses such as tissue repair, neuroprotection, and immunosuppression [29–32]. Therefore, we propose that GA inhibits the fibrosis process by targeting the SUMOylation modification pathway of macrophages and regulating their immunophenotype.

The molecular mechanism by which SUMOylation drives the fibrotic process by targeting and regulating the core components of the TGF-β/Smad pathway (SUMOylation of TGFβRⅠ enhances the phosphorylation of Smad2/3, SUMOylation of Smad2 increases its phosphorylation level and transcriptional activity, and SUMOylation of Smad4 mediates ECM remodeling) has been clarified [33–35]. GA is a negative regulator of the SUMOylation modification of Smad4. In our experiment, *E.granulosus* infection significantly enhanced the SUMOylation modification of Smad4, increased the level of serum TGF-β1, and exacerbated fibrosis. GA intervention not only significantly inhibits the SUMOylation modification of Smad4, but also hinders the nuclear translocation process of SUMOylation Smad4 in macrophages induced by EgCF (verified by the nuclear-cytoplasmic separation experiment), thereby reversing the M2-type polarization phenotype (CD206↓). And synergistically inhibit the proliferation and activation of HSCs (α-SMA / PCNA↓). To sum up, the SUMO modification of Smad4 may be the key factor for GA to inhibit macrophage polarization and resist echinococcosis fibrosis.

This study reveals for the first time that SUMOylation of Smad4 regulates the M2 polarization of macrophages and the crosstalk between macrophages and HSCs, thereby promoting the progression of hepatic fibrosis mediated by *E.granulosus* infection. This finding not only provides an important theoretical basis for a deeper understanding of the pathogenesis of parasitic liver fibrosis, but also proposes Smad4 SUMOylation as a new target for intervention with great potential for anti-fibrosis, and finds that GA as an effective inhibitor of the target can significantly inhibit the above pathogenic pathways and exert anti-fibrosis effects. It provides innovative strategies and drug candidates for anti-fibrotic treatment of hydatid disease. Of course, this study still has some limitations. Firstly, the *in vivo* pharmacokinetics and long-term safety of GA have not been systematically evaluated. Secondly, the specific inhibition of Smad4 SUMOylation by GA needs to be further verified. In addition, the core factors mediating the pathogenic crosstalk between macrophages and HSCs still need to be screened and identified. These will provide key directions for subsequent targeted therapy research.

### Conclusions

Taken together, the present study clarified that the imbalance of SUMOylation is a central molecular hub in the progression of liver fibrosis induced by *E. granulosus* infection. GA can specifically inhibit the SUMOylation of Smad4 and its nuclear translocation by targeting this pathway, thereby effectively reversing the M2 macrophage over-polarization and inhibiting the activation and proliferation of hepatic stellate cells. More importantly, GA significantly disrupted the vicious cycle of mutual promotion between M2-type macrophages and activated HSCs. Therefore, GA exerts its anti-hydatid liver fibrosis effect through multi-target intervention of SUMOylation - Smad4 signaling axis and cell-cell interaction network, which provides an important experimental basis for targeted therapy based on SUMOylation regulation.

## Authorship contribution

Xiangwei Wu and Xueling Chen conceived the study. Qiuyue Chen, Xueting Yu, Xinyu Jiang: Data curation, Visualization, Writing – original draft. Huijiao Jiang, Jun Hou, Lianghai Wang and Junying Xu: Methodology, Investigation, Formal analysis. Dan Dong and Qiuyue Chen: Writing – review & editing. Xiangwei Wu, Xueling Chen, Dan Dong and Huijiao Jiang: Funding acquisition. Xiangwei Wu and Xueling Chen: Conceptualization, Supervision. All authors revised the manuscript critically for important intellectual content and approved the final version.

## Acknowledgements

The study was approved by the National Key R&D Program of China (2024YFC2309700); Tianshan Young Talent Scientific and Technological Innovation Team (2023TSYCTD0020); Corps Guidance Science and Technology Plan Project (2023ZD034, 2024ZD018); Shihezi University Scientific Research Project (ZZZC2023026).

---

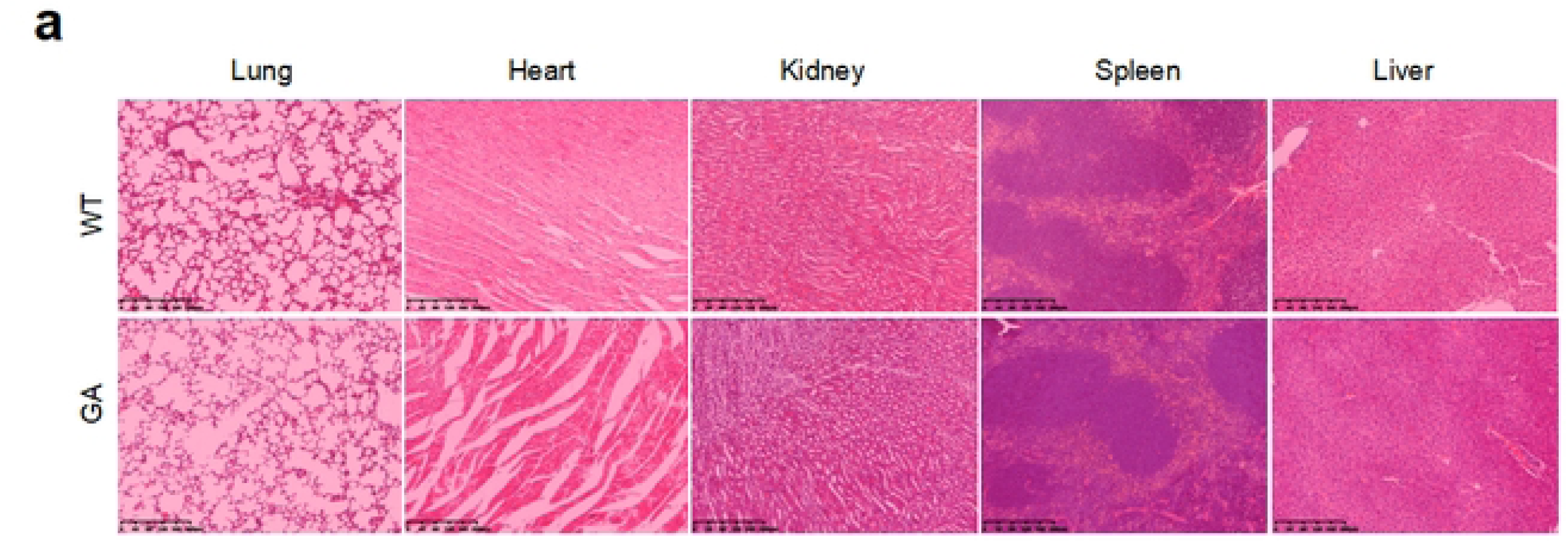

